# Does the Preferred Walk-Run Transition Speed on Inclines Minimize Energetic Cost, Heart Rate or Neither?

**DOI:** 10.1101/2020.07.11.198796

**Authors:** Jackson W. Brill, Rodger Kram

## Abstract

Humans prefer to walk at slow speeds and to run at fast speeds. In between, there is a speed at which people choose to transition between gaits, the Preferred Transition Speed (PTS). At slow speeds, it is energetically cheaper to walk and at faster speeds, it is cheaper to run. Thus, there is an intermediate speed, the Energetically Optimal Transition Speed (EOTS). Our goals were to determine: 1) how PTS and EOTS compare across a wide range of inclines and 2) if the EOTS can be predicted by the heart rate optimal transition speed (HROTS). Ten healthy, high-caliber, male trail/mountain runners participated. On day 1, subjects completed 0° and 15° trials and on day 2, 5° and 10°. We calculated PTS as the average of the walk-to-run transition speed (WRTS) and the run-to-walk transition speed (RWTS) determined with an incremental protocol. We calculated EOTS and HROTS from energetic cost and heart rate data for walking and running near the expected EOTS for each incline. The intersection of the walking and running linear regression equations defined EOTS and HROTS. We found that PTS, EOTS, and HROTS all were slower on steeper inclines. PTS was slower than EOTS at 0°, 5°, and 10°, but the two converged at 15°. PTS and EOTS were only moderately correlated. Although EOTS correlated with HROTS, EOTS was not predicted accurately by heart rate on an individual basis.

## INTRODUCTION

Humans prefer to walk at slow speeds and to run at fast speeds. In between, there is a speed at which people choose to transition between gaits, the Preferred Transition Speed (PTS). At slow speeds, it is energetically cheaper to walk and at faster speeds, it is cheaper to run. Thus, there is an intermediate speed, the Energetically Optimal Transition Speed (EOTS). The prevailing scientific thinking prior to the 1990s was that at a given speed, humans and other species choose the gait that minimizes energy expenditure, i.e. that PTS = EOTS. But more recent research indicates that the PTS occurs at a speed slower than the EOTS in humans (Abe et. al. 2019, Ganley et. al. 2011, Hreljac 1993, Minetti et. al. 1994, Rotstein et. al. 2005). Curiously, in horses the walk-trot transition occurs near the EOTS (Griffin et. al. 2004) but the trot-gallop transition is triggered by mechanics, not energetics (Farley and Taylor 1991).

Consequently, researchers have proposed various neuromechanical hypotheses to explain the human walk-run gait transition speed, but no clear consensus has emerged (Kung et. al. 2018). Several researchers have hypothesized that the walk-run transition is triggered by impending fatigue of specific muscles, notably the tibialis anterior (Abe et. al. 2019, Bartlett and Kram 2008, Hreljac 1995, Hreljac et. al. 2001, Hreljac et. al. 2008). Neptune and Sasaki (2005) implicated force insufficiency due to the muscle force-length and force-velocity relationships as another walk-run gait transition trigger. They found that when walking at the PTS, the plantar flexor muscle force production was insufficient, despite an increase in EMG muscle activation. The physics of the body’s center of mass inverted pendulum-like motion is another proposed trigger (Kram et al. 1997, Usherwood 2005).

The PTS is slower on inclines relative to flat terrain (Diedrich and Warren 1998, Hreljac 1995, Hubel and Usherwood 2013, Minetti et. al. 1994). Further, the PTS is slower than the EOTS on inclines up to 8.5° and the difference between the PTS and EOTS remains fairly constant (∼0.2 m/s) on those moderate grades (Minetti et al. 1994). Inclined locomotion is a promising tool for scientists trying to understand what triggers the walk-run transition in general (Hreljac 1995) because compared to level locomotion, uphill locomotion has a greater energetic cost and places different demands on specific muscle groups (Vernillo et. al. 2016).

On an applied note, the walk-run gait transition on inclines is of great interest to competitive trail/mountain runners who often ponder whether they should walk or run when racing uphill. It is obvious that competitors should nearly always run during flat and downhill sections of any race. But, dynamic factors such as the steepness of the incline, speed, ground surface, length of the climb, and the overall duration/length of the race all add complexity to an individual’s gait selection process. Many trail/mountain runners use GPS watches and heart rate monitors to guide their training and racing (Koop and Rutberg, 2016). If heart rate is an accurate proxy for energetic cost, heart rate monitors combined with GPS sensors may allow athletes to identify their EOTS during a race or in training, choose their gait accordingly, and thus enhance performance.

The purpose of this study was to determine how the PTS and EOTS change over a wide range of inclines. Specifically, we investigated how incline affects the PTS, EOTS and the relationship between the two. We also determined how heart rate is influenced by gait selection. Our first hypothesis was that PTS and EOTS would both be slower on steeper inclines. We also tested the null hypothesis that the absolute difference in speed between the PTS and EOTS would be independent of incline. We anticipated rejecting that hypothesis because we thought that with the greater energetic demand on steeper inclines, there would be a greater drive to minimize energetic cost. That is, we expected that PTS and EOTS would converge on steeper inclines. Finally, we hypothesized that the EOTS would equal the heart rate optimal transition speed (HROTS) at each incline. We thought this would occur because energetic cost generally correlates with heart rate during steady-state endurance exercise (Arts and Kuipers 1994).

## METHODS

### Subjects

Ten healthy, high-caliber, male trail/mountain runners (28.7 ± 5.7 yr, 67.6 ± 4.9 kg 1.79 ± 0.06 m; mean ±SD) volunteered and provided informed consent as per the University of Colorado Institutional Review Board. All subjects had placed in the top 10% in a trail/mountain running competition within the previous two years.

### Experimental design

The study consisted of two sessions. On Day 1, we collected data for walking and running at 0° and then at 15° (26.8% grade). On Day 2, we collected data for walking and running at 5° (8.7% grade) and then 10° (17.6% grade). We did not randomize the trial order to avoid having a subject complete the more difficult 10° and 15° trials back-to-back on the same day. For each incline, we first determined PTS and then collected energetics and heart rate data simultaneously to calculate EOTS and HROTS. For each subject, we randomly assigned half the subjects to the “walk-first” gait order and half to the “run-first” gait order. Subjects walked and ran on a classic Quinton 18-60 motorized treadmill with a rigid steel deck (Quinton Instrument Company, Bothell, WA).

### Determination of PTS

The average of the walk-to-run transition speed (WRTS) and run-to-walk transition speed (RWTS) defined the PTS as per Hreljac et. al. (2007). We first determined the WRTS in the walk-first group and then their RWTS and *vice versa* for the run-first group. Based on pilot experiments, we selected starting speeds such that there was no doubt which gait would be preferred at the initial speed. Once the speed of the treadmill was correctly set, subjects mounted the treadmill and chose their gait *ad libitum*. After we determined the preferred gait at the particular speed, the subject straddled the treadmill belt while we changed the speed by 0.1 m/s (increased during WRTS trials, decreased during RWTS trials). The process repeated until a gait transition occurred and was sustained for 30 seconds.

### Determination of EOTS and HROTS

For the energetics and heart rate trials, we set the initial speed based on pilot experiments that indicated it would be near the EOTS. Subjects in the walk-first group walked at the incline-specific initial speed for 5 min, rested for ∼5 min and then ran at that speed for 5 min. Subjects in the run-first group did the opposite. During the rest periods, we re-weighed the subject and they drank just enough water to compensate for the weight loss due mostly to sweating. Thus, each subject maintained a nearly constant weight throughout all the trials.

To measure metabolic rate during walking and running, we used an open-circuit, expired gas analysis system (TrueOne 2400; ParvoMedics, Sandy, UT). Subjects wore a mouthpiece with a one-way breathing valve and a nose clip allowing us to collect their expired air. The ParvoMedics software calculated the STPD rates of oxygen consumption (V□O_2_) and carbon dioxide production (V□CO_2_) and we averaged the last 2 minutes of each 5-minute trial. We then calculated metabolic power using the equation of Péronnet and Massicotte (1991) equation, as clarified by Kipp et al. (2018). We only included trials with respiratory exchange ratios (RER) <1.0 to ensure that metabolic energy was predominantly being provided from oxidative pathways. We used an R7 Polar iWL (Polar Electro Oy, Kempele, Finland) to measure heart rate in beats per minute (bpm) and averaged the values for the last 2 min of each trial.

Immediately after both gait trials were completed for the initial speed, we calculated and compared the metabolic power required for walking and running. If walking was the more economical gait, we increased the treadmill speed by 0.1 m/s, and the process repeated. If running was the more economical gait, we decreased the treadmill speed by 0.1 m/s, and the process repeated. Each subject performed three speeds, both walking and running at each incline. However, some subjects needed to complete walking and running trials at a fourth speed so that we could obtain energetics data for one speed faster and one speed slower than their EOTS.

### Data Analysis

We used R Studio (https://rstudio.com/) for all statistical analysis. For the three speeds at which the differences between metabolic rates between walking and running were least, we calculated linear regression equations for both metabolic power and heart rate as functions of speed for both walking and running for each subject and incline. The speeds at which the two equations intersected defined the EOTS and HROTS for each subject.

Overall, we analyzed ten subjects at four different inclines, i.e. 40 determinations of EOTS and HROTS. Of those 80 linear regression analyses, the walking vs. running regressions intersected at a speed < 3 m/sec for all but two subjects (one subject for EOTS at 15° and a different subject for HROTS at 10°). Essentially, those individuals’ regression lines were nearly parallel. We chose to exclude those two conditions from further statistical analysis and aggregate data compilation. We then determined the linear regression equations and R^2^ values for PTS, EOTS, and HROTS as functions of incline. Furthermore, we compared PTS vs. EOTS, PTS vs. HROTS, and EOTS vs. HROTS at each incline by computing p-values (from paired t-tests), R^2^ values (from linear regression analysis), and effect sizes (from Cohen’s d statistic).

## RESULTS

As expected, on the level and at all inclines, walking generally required less metabolic power at slow speeds and running required less at faster speeds. Thus, regression lines for metabolic power vs. speed in the two gaits generally intersected. An example for one subject at 15° is depicted in Figure 1A. Heart rates also generally showed similar patterns and an example of the heart rate optimal transition speed (HROTS) is shown in Figure 1B.

**Figure 1.**
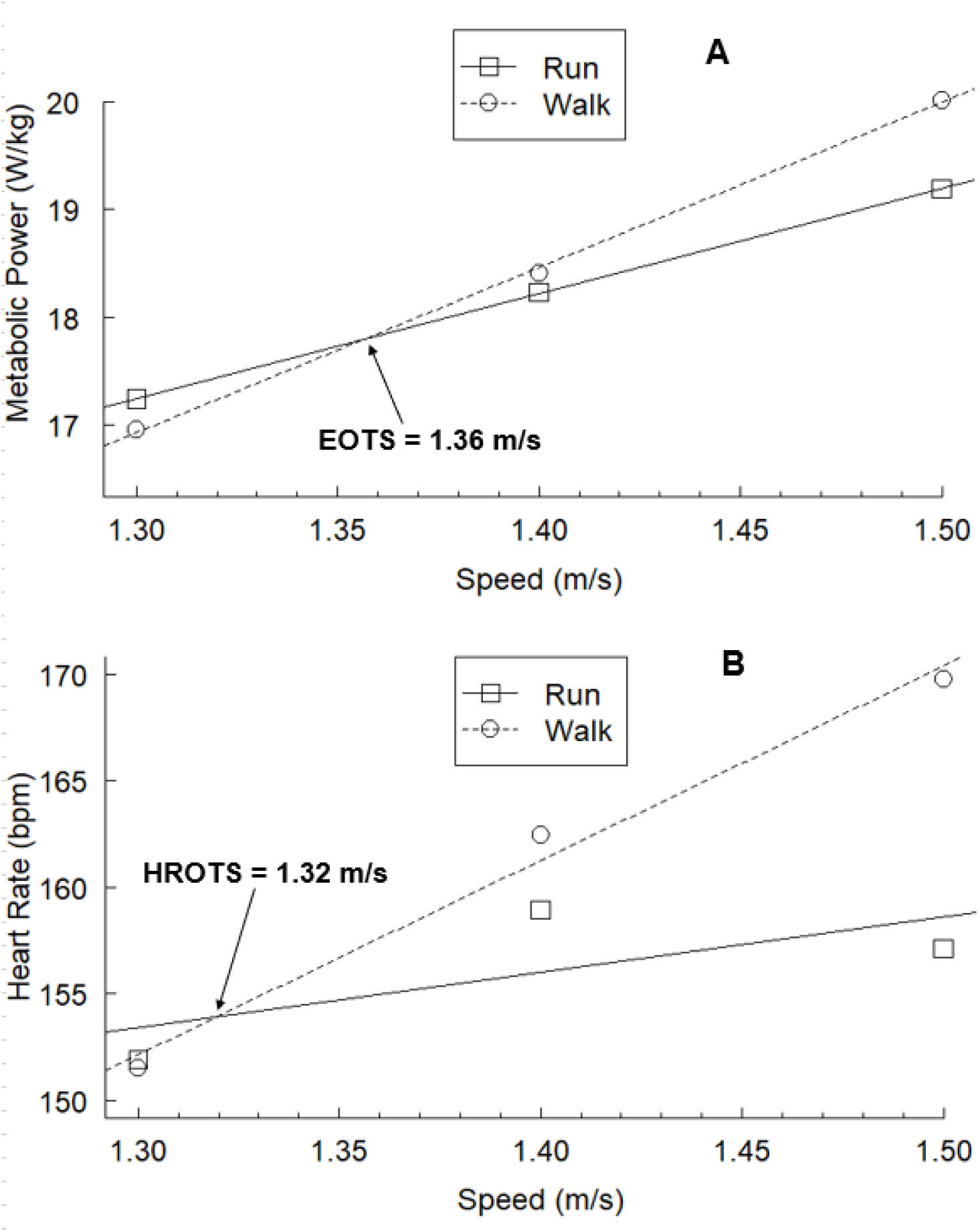
A. An example of an energetically optimal transition speed (EOTS) linear regression analysis for one subject on a 15° incline. B. An example of a heart rate optimal transition speed (HROTS) linear regression analysis for the same subject on a 15° incline. Heart rate was measured in beats per minute (bpm).

### Preferred Transition Speed

PTS was slower on steeper inclines (Table 1, Figure 2A). The linear regression equation for the PTS (in m/s) vs. incline (Θ, in degrees) was:

**Table 1:** Preferred (PTS), energetically optimal (EOTS), and heart rate optimal (HROTS) walk-run transition speed averages for each incline. Results presented as mean ± SD. N=10 for all values except for EOTS at 15° (n=9) and HROTS at 10° (n=9).

| **Incline (°)** | **0** | **5** | **10** | **15** |
| --- | --- | --- | --- | --- |
| **PTS (m/s)** | 1.95 ± 0.12 | 1.78 ± 0.27 | 1.62 ± 0.14 | 1.47 ± 0.21 |
| **EOTS (m/s)** | 2.14 ± 0.10 | 1.99 ± 0.13 | 1.78 ± 0.13 | 1.51 ± 0.20 |
| **HROTS (m/s)** | 2.13 ± 0.14 | 1.89 ± 0.13 | 1.69 ± 0.23 | 1.46 ± 0.24 |

**Figure 2.**
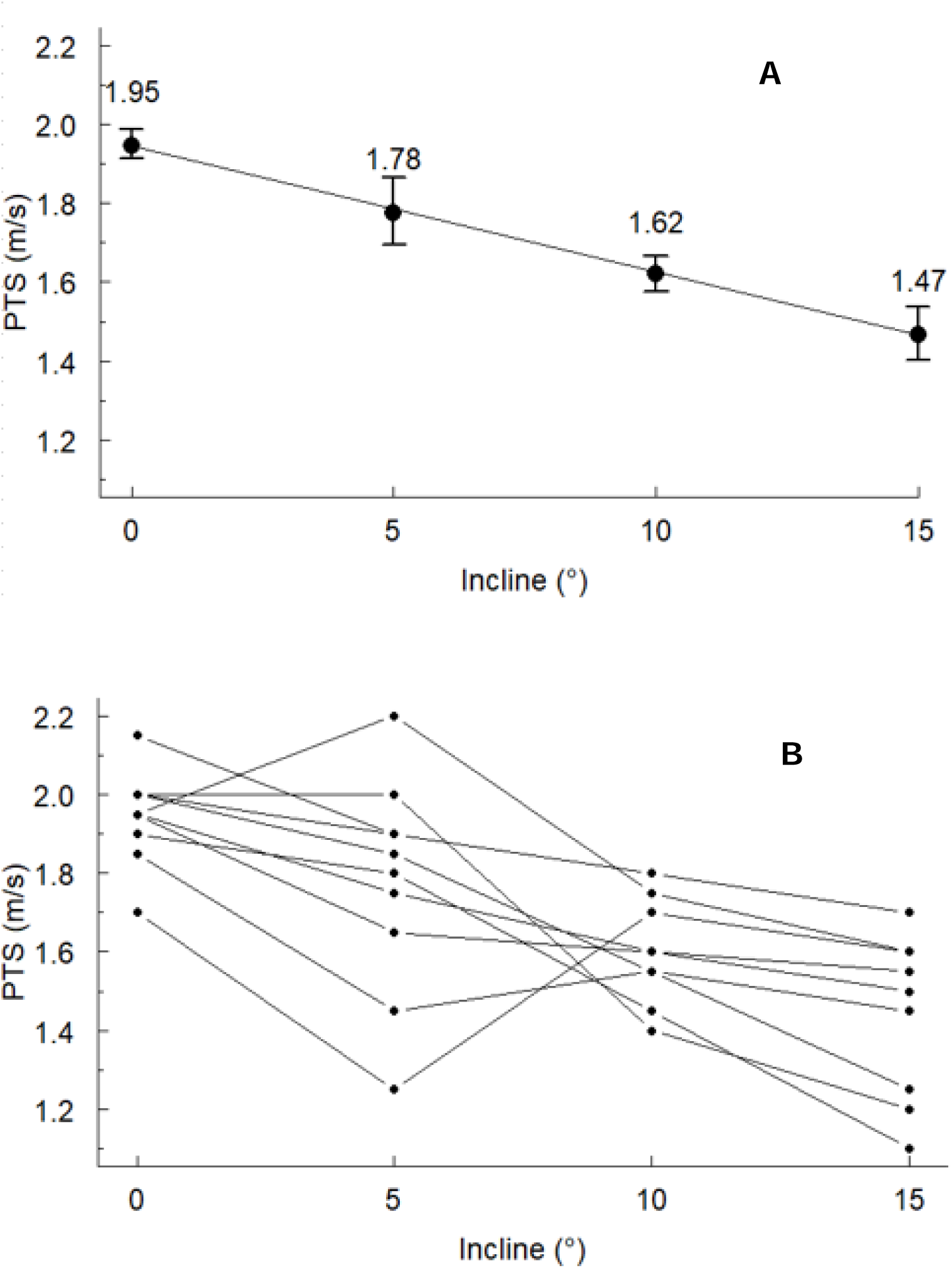
A. Preferred Transition Speed (PTS) vs. incline. Symbols are means +/- SEM. The overall linear regression equation and R^2^ value were calculated from 4 inclines and n=10 subjects (40 total data points). B. PTS data for each subject presented individually.

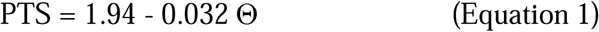

The slope of the regression was significantly different from zero (p=6.90 e-7 and R^2^=0.481). Seven of the subjects’ PTS decreased monotonically with incline while three subjects had slight deviations from that overall pattern (Figure 2B).

### Energetically Optimal Transition Speed

EOTS was slower on steeper inclines (Table 1, Figure 3A). The linear regression equation for the EOTS (in m/s) vs. incline (Θ, in degrees) was:

**Figure 3.**
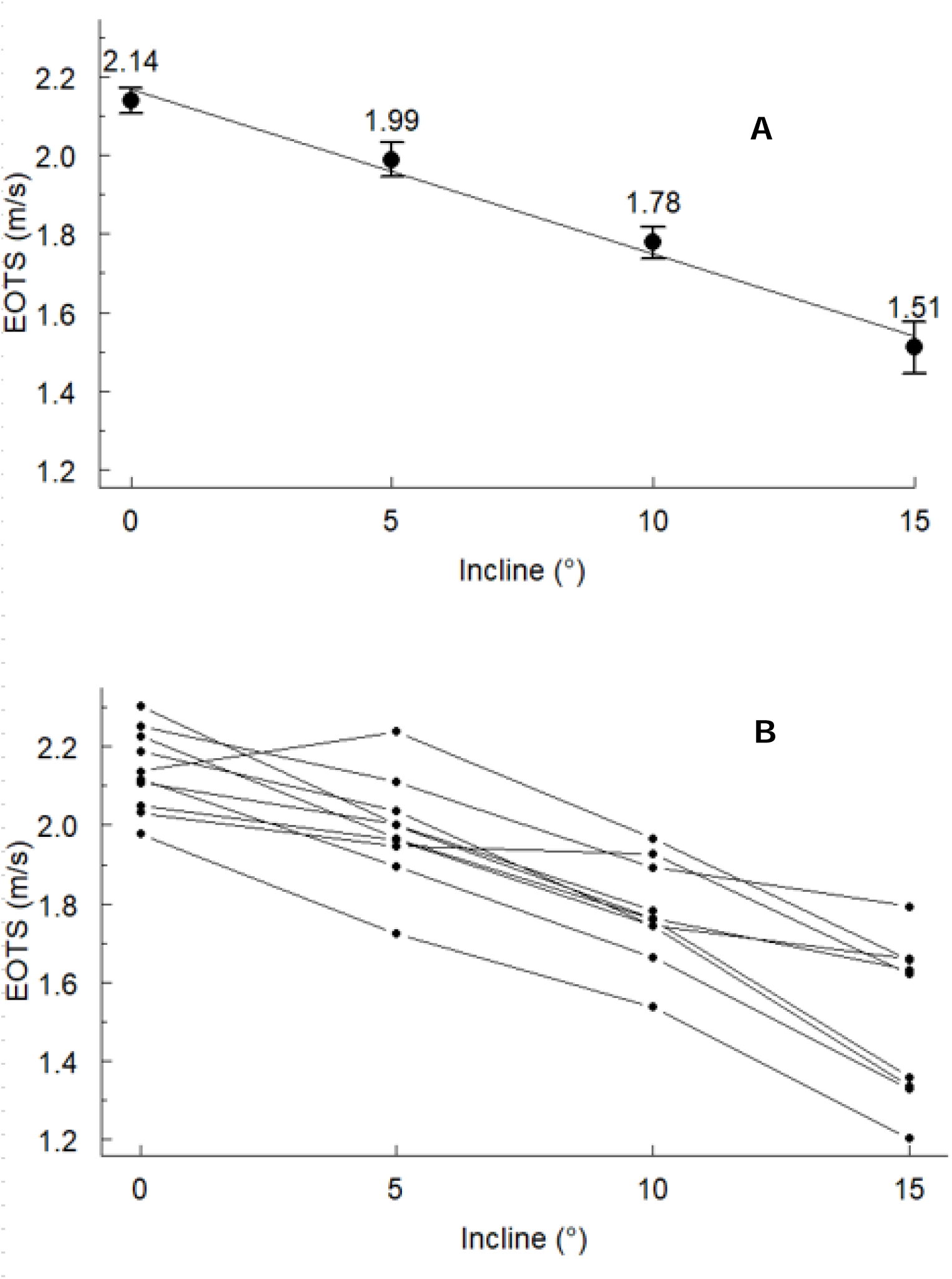
A. Energetically Optimal Transition Speed (EOTS) vs. incline. Symbols are means +/- SEM. The overall linear regression equation and R^2^ value were calculated from 4 inclines and n=10 subjects at 0°, 5°, and 10°, but n=9 subjects at 15° (39 total data points). B. EOTS data for each subject presented individually.

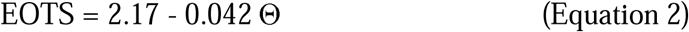

The slope of the regression was significantly different from zero (p=4.22 e-12 and R^2^=0.731). Eight of the subjects’ EOTS decreased monotonically with incline while two subjects had slight deviations from that overall pattern (Figure 3B).

### Heart Rate Optimal Transition Speed

HROTS was slower on steeper inclines (Table 1, Figure 4A). The linear regression equation for the HROTS (in m/s) vs. incline (Θ, in degrees) was:

**Figure 4.**
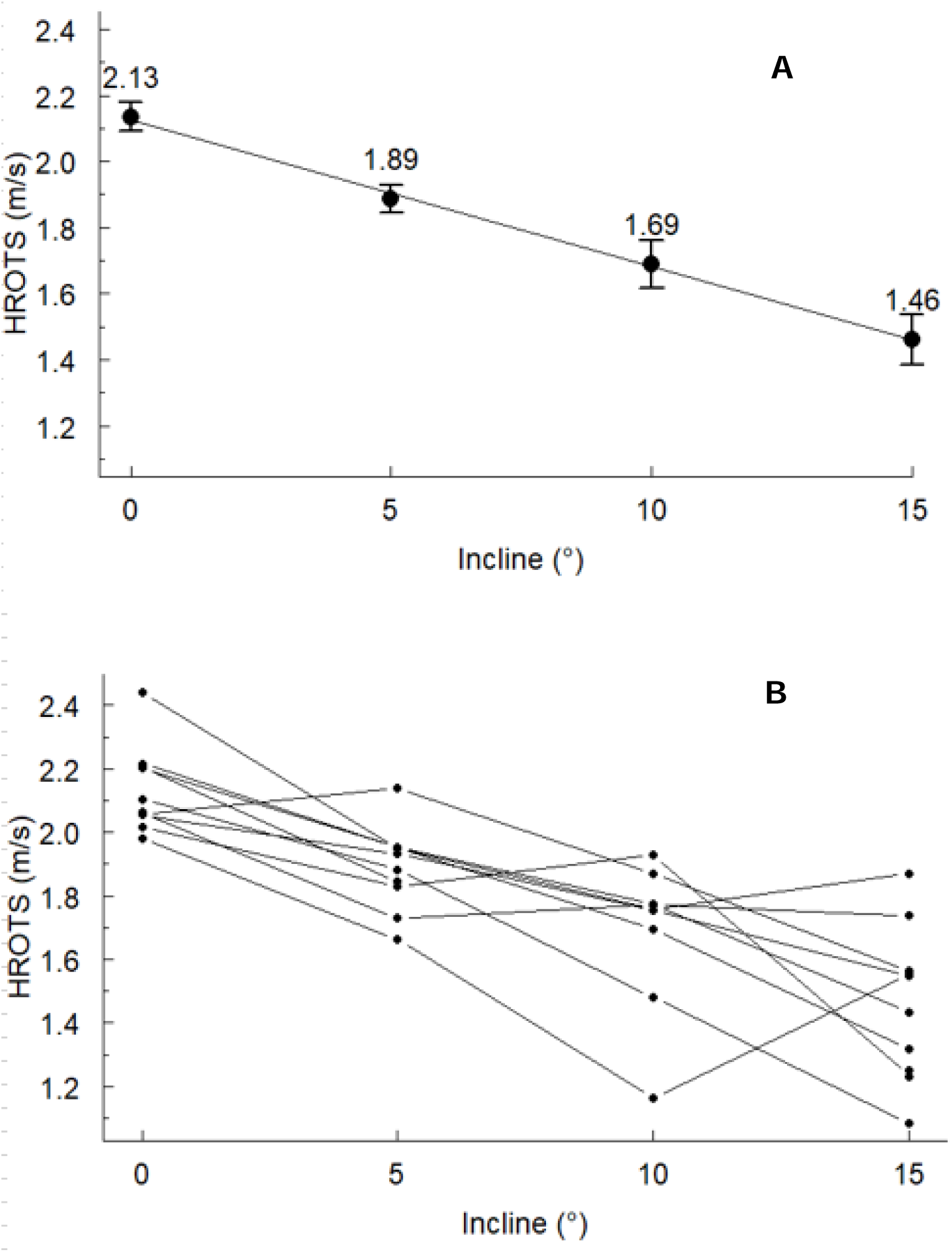
A. Heart Rate Optimal Transition Speed (HROTS) vs. incline. Symbols are means +/- SEM. The overall linear regression equation and R^2^ value were calculated from 4 inclines and n=10 subjects at 0°, 5°, and 15°, but n=9 subjects at 10° (39 total data points). B. HROTS data for each subject presented individually.

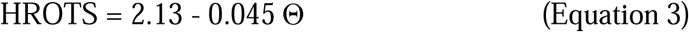

The slope of the regression was significantly different from zero (p= 4.56 e-10 and R^2^=0.655). Among individuals, HROTS generally decreased but was more variable than the other transition speed metrics (Figure 4B).

### Comparisons between PTS, EOTS, and HROTS at each incline

Mean PTS was slower than mean EOTS at 0°, 5°, and 10°, but not at 15° (p-values of 0.002, 0.107, 0.007, and 0.931, respectively; Cohen’s d effect sizes of 1.77, 1.04, 1.17, and 0.02, respectively). The correlations between PTS and EOTS were weak to moderate at all inclines (R^2^ values of 0.045, 0.478, 0.193, and 0.498 at 0°, 5°, 10°, and 15°, respectively; p-values comparing these slopes to zero were 0.555, 0.027, 0.204, and 0.034, respectively).

Mean PTS was slower than mean HROTS at 0°, but not at 5°, 10°, and 15° (p-values of 0.001, 0.366, 0.271, and 0.946, respectively; Cohen’s d effect sizes of 1.48, 0.54, 0.44, and 0.03, respectively). The correlations between PTS and HROTS were weak to moderate at all inclines (R^2^ values of 0.270, 0.371, 0.133, and 0.002 at 0°, 5°, 10°, and 15°, respectively; p-values comparing these slopes to zero were 0.124, 0.062, 0.335, and 0.898, respectively).

The relationship between EOTS and HROTS was inconsistent across inclines (Figure 5). Mean EOTS was not different than mean HROTS at 0°, was faster than mean HROTS at 5°, was similar at 10°, and was not different at 15° (p-values of 0.818, 0.002, 0.124 and 0.503, respectively; Cohen’s d effect sizes of 0.05, 0.77, 0.40, and 0.21, respectively). The correlations between EOTS and HROTS were moderate to strong at all inclines (R^2^ values of 0.658, 0.708, 0.567, and 0.376 at 0°, 5°, 10°, and 15°, respectively; p-values comparing these slopes to zero were 0.004, 0.002, 0.019, and 0.079, respectively).

**Figure 5.**
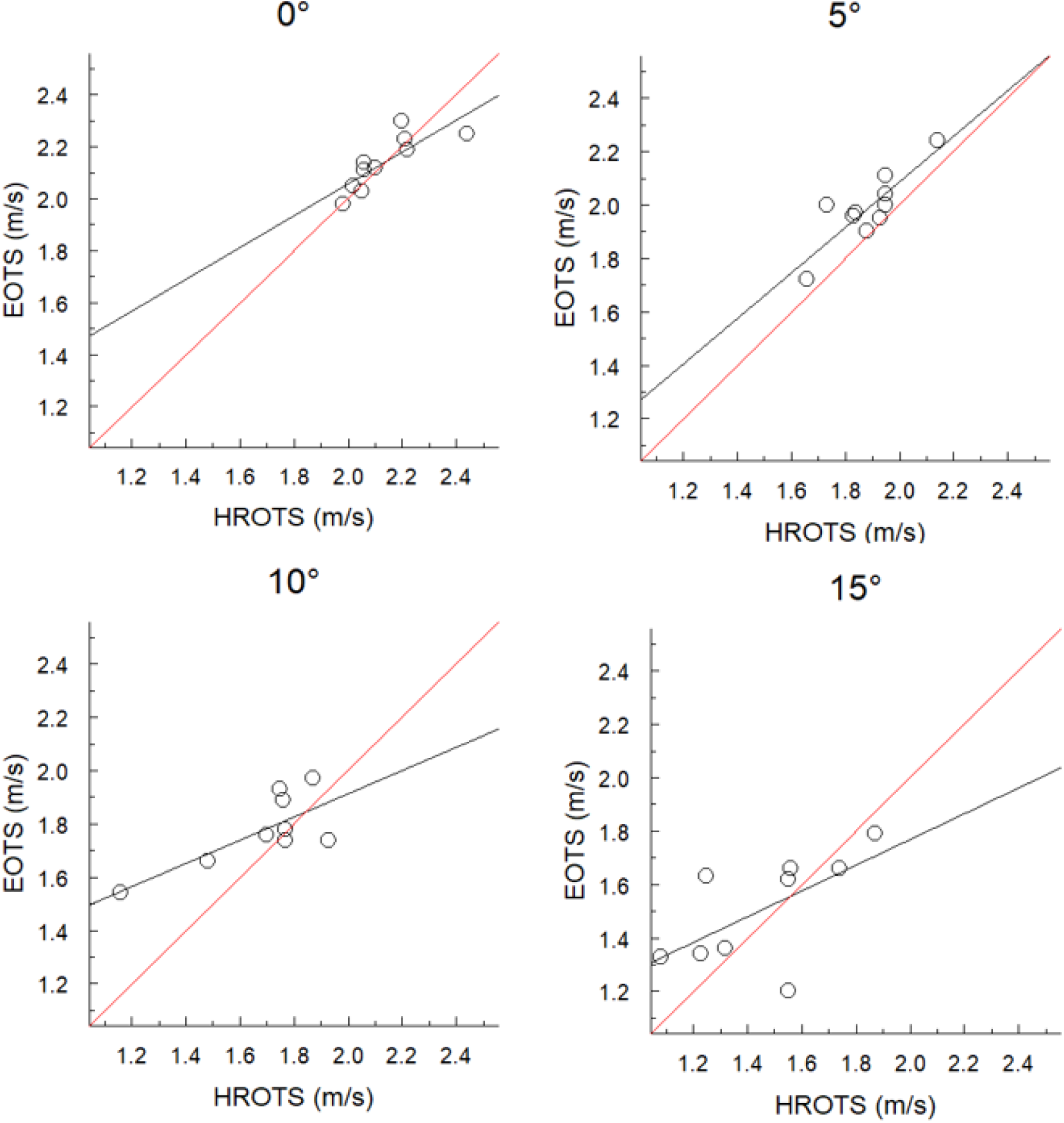
Energetic (EOTS) vs. heart rate (HROTS) optimal transition speeds for treadmill inclination angles of 0° (n=10), 5° (n=10), 10° (n=9), and 15° (n=9). The regression is shown by the black line, while the red line is the line of identity.

## DISCUSSION

We retain our first hypothesis that PTS and EOTS would both be slower on steeper inclines. This is consistent with previous research. Minetti et. al. (1994) determined that both PTS and EOTS were slower on steeper inclines up to 8.5°. Diedrich and Warren (1998), Hreljac (1995), and Hubel and Usherwood (2013) all determined that PTS was slower on inclines (but did not measure EOTS).

Hubel and Usherwood (2013) also developed a prediction for the walk-run transition on inclines using the physics-based limits of an inverted pendulum/compass gait model for walking. Their prediction reduces to a simple rule of thumb whereby the PTS slows by about 1% for every 1% increase in incline (or ∼1.8% per degree). In the present study, the mean PTSs at 5°, 10°, and 15° were 1.78, 1.62, and 1.47 m/s, respectively. Using Hubel and Usherwood’s (2013) rule of thumb, with the baseline PTS at 0° found in the present study (1.95 m/s), the predicted PTSs at 5°, 10°, and 15° are 1.78, 1.63, and 1.49 m/s, respectively. Our empirical measurements coincide remarkably well with the predicted values. This supports Hubel and Usherwood’s (2013) conclusion that the inverted pendulum/compass gait model for walking informs the qualitative understanding of why PTS slows with incline and also has predictive ability in quantifying PTS across inclines. However, on an individual level, the predictive ability was less accurate. Using Hubel and Usherwood’s (2013) rule of thumb, with the baseline PTS at 0° for each subject used to predict their PTS at 5°, 10°, and 15°, the average absolute percent differences between predicted and actual PTS were 8.8%, 7.1%, and 11.7% at 5°, 10°, and 15°, respectively.

We also retain our second hypothesis that PTS and EOTS would converge at steeper inclines. The difference between the average PTS and EOTS was 0.19 m/s at 0°, 0.21 m/s at 5°, 0.16 m/s at 10°, and 0.04 m/s at 15°, with PTS always numerically slower than EOTS. Statistical analysis indicated that the PTS was slower than EOTS at 0°, 5°, and 10° but not at 15°. The mean absolute difference between the PTS and EOTS at 5° was numerically greater than any other incline but had inexplicably large inter-subject variability (standard deviation at 5° was approximately twice that at 0° and 10°).

The finding that PTS was slower than EOTS on level terrain was previously well established (Abe et. al. 2019, Ganley et. al. 2011, Hreljac 1993, Minetti et. al. 1994, Rotstein et. al. 2005), with just one study finding equal PTS and EOTS (Mercier et. al. 1994). Only Minetti et. al. (1994) have previously studied how the relationship between PTS and EOTS is affected by incline. They concluded that the absolute difference (m/s) between PTS and EOTS does not change with gradient, but they studied more moderate inclines, up to 8.5°.

Our data demonstrate that the difference between PTS and EOTS remained roughly constant up to 10°, but not at 15°. Clearly, optimizing energetic economy does not solely determine PTS at moderate grades that do not involve a high rate of exertion. As grade steepens, energetic cost may become more influential. However, the equation for the regression of PTS vs. EOTS at 15° (Figure 7) has an R^2^ value of only 0.498, i.e., PTS only explained half of the variance in EOTS. Despite the fact that PTS and EOTS did converge at 15°, further investigation on the relationship between PTS and EOTS at even steeper inclines would be interesting.

**Figure 6.**
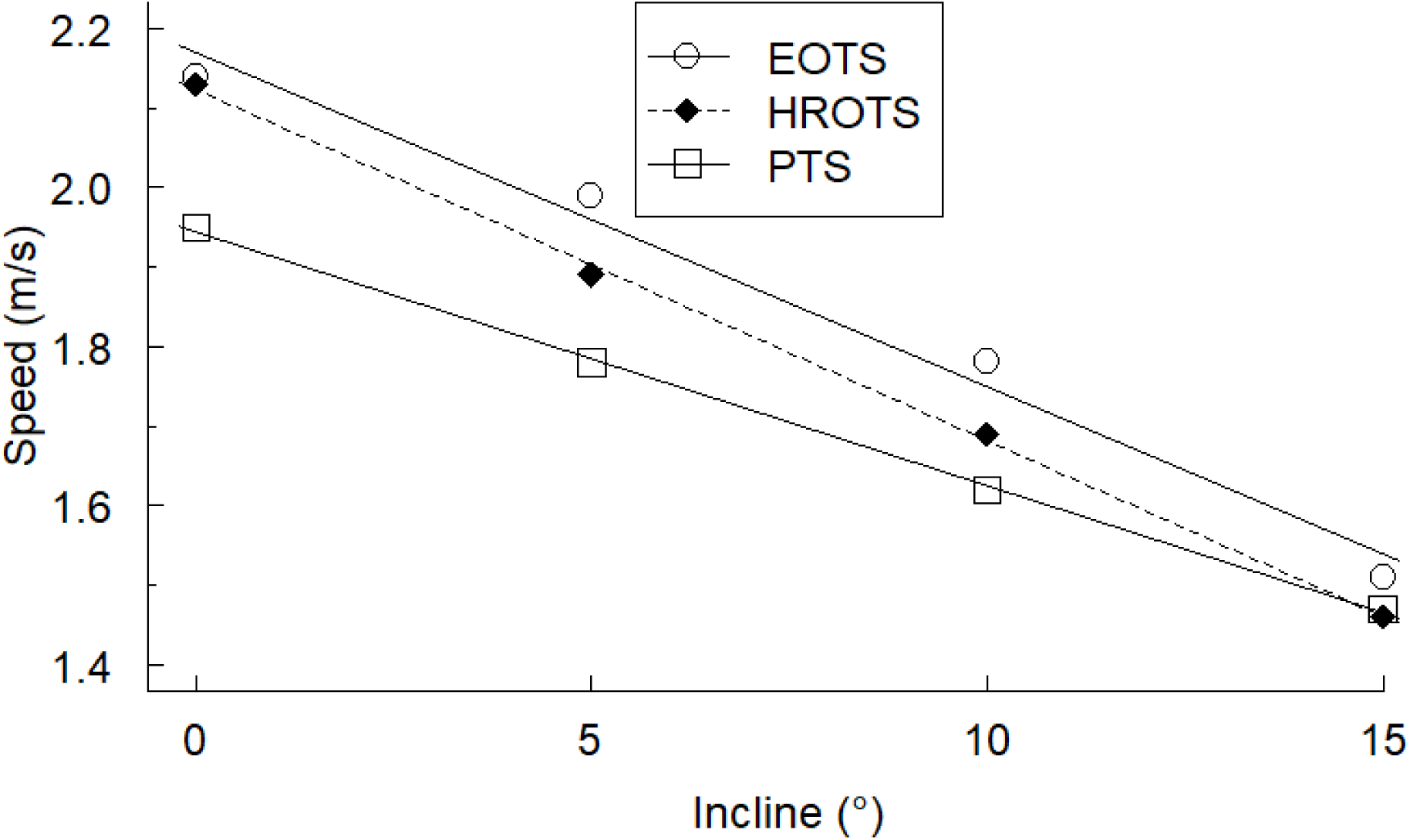
Mean preferred (PTS), energetically optimal (EOTS), and heart rate optimal (HROTS) transition speeds for each incline.

**Figure 7:**
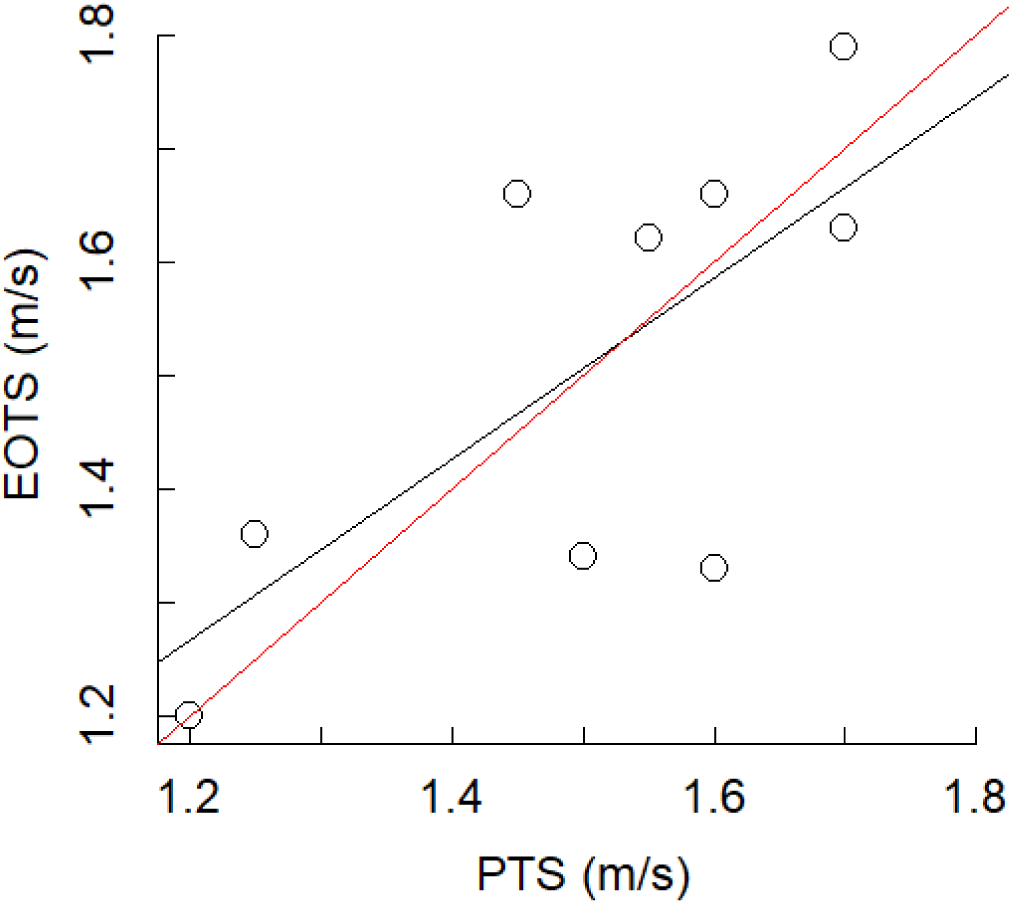
PTS vs. EOTS data (n=9) at 15°. The regression is shown by the black line, while the red line is the line of identity. The R^2^ value of the regression is 0.498 with p-value = 0.034 (compared to a slope of 0).

The relationship between PTS and EOTS, and thus whether energetic cost serves as a trigger for gait transitions, has been studied in horses as well. Like humans, the evidence for PTS equaling EOTS in these quadrupeds is nuanced. Hoyt and Taylor (1981) concluded that at their preferred speed within each gait, horses choose the gait that minimizes energetic cost. Griffin et. al. (2004) built upon that idea and found that energetics also seem to trigger the walk-trot transition. But Farley and Taylor (1991) clearly demonstrated that the trot-gallop transition was triggered by musculoskeletal forces and not energetics. In humans, there is also some evidence for kinetic factors triggering the level walk-run transition (Raynor et. al. 2002, Hreljac et. al. 2008).

Our third hypothesis was that EOTS would equal HROTS at all inclines. Statistical analysis indicated that the EOTS and HROTS were not different at 0°, diverged at 5°, were similar at 10°, and were again not different at 15°. Furthermore, the correlations between EOTS and HROTS were not consistent enough to have practical predictive application for competitive trail/mountain runners (Figure 5). Thus, we reject our third hypothesis for all inclines besides 0°. Rotstein et. al. (2005) and Mercier et. al. (1994) found that PTS equals HROTS on level terrain, although Mercier et. al. (1994) also found that PTS equals EOTS, which differs from the rest of the research done on this topic. At 0°, we found that EOTS and HROTS were equal and both were faster than PTS (Table 2, Figure 6). No previous research has studied HROTS on inclines.

**Table 2:**
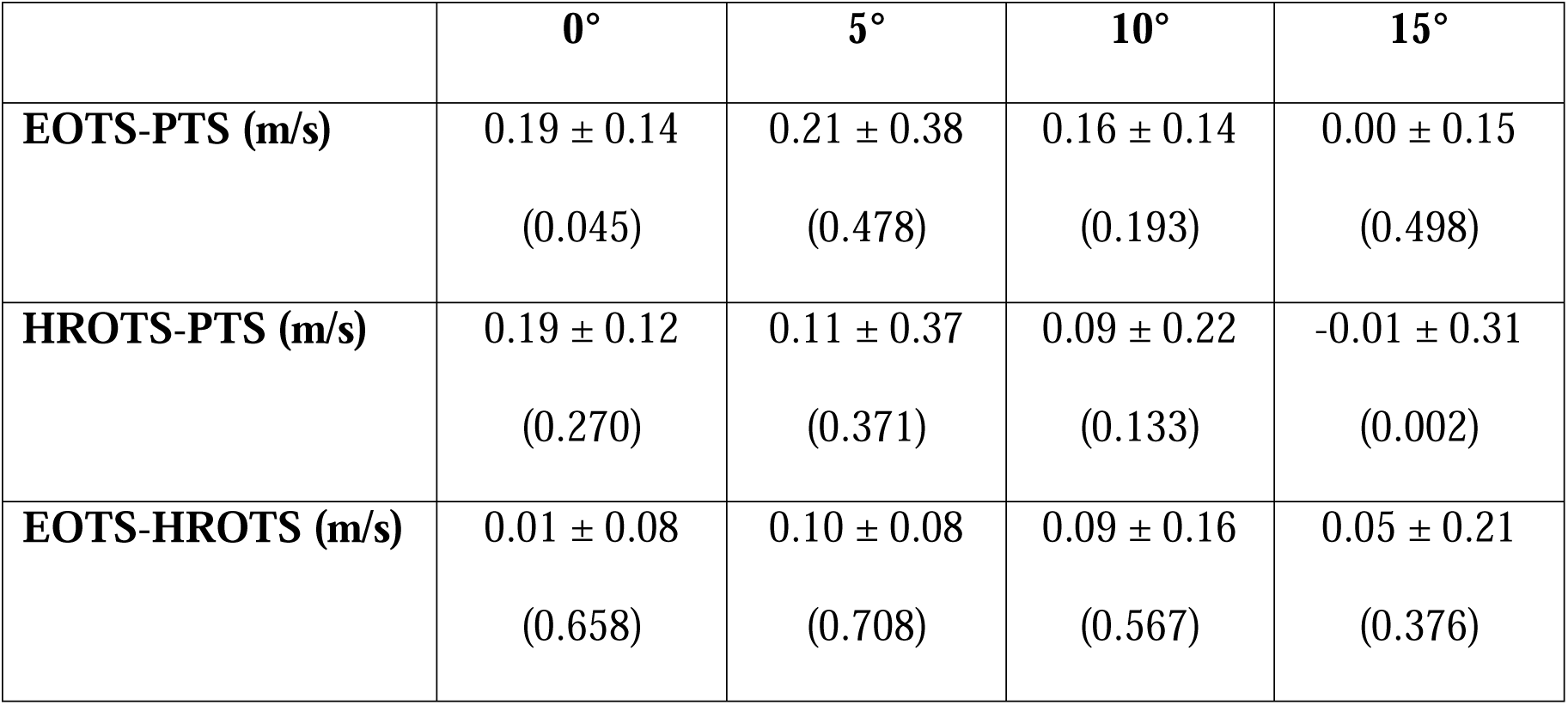
Average differences and coefficient of determinations (R^2^) among subjects in the transition speeds for each incline. Results presented as mean ± SD (R^2^). Differences for each subject were averaged, as opposed to taking the difference of the average PTS, EOTS, and HROTS.

|  | **0°** | **5°** | **10°** | **15°** |
| --- | --- | --- | --- | --- |
| **EOTS-PTS (m/s)** | 0.19 ± 0.14<br>(0.045) | 0.21 ± 0.38<br>(0.478) | 0.16 ± 0.14<br>(0.193) | 0.00 ± 0.15<br>(0.498) |
| **HROTS-PTS (m/s)** | 0.19 ± 0.12<br>(0.270) | 0.11 ± 0.37<br>(0.371) | 0.09 ± 0.22<br>(0.133) | -0.01 ± 0.31<br>(0.002) |
| **EOTS-HROTS (m/s)** | 0.01 ± 0.08<br>(0.658) | 0.10 ± 0.08<br>(0.708) | 0.09 ± 0.16<br>(0.567) | 0.05 ± 0.21<br>(0.376) |

Our lab’s previous research on steep uphill running and walking has focused on 30° (Giovanelli et. al. 2016, Ortiz et. al. 2017) because it is the optimal incline for vertical kilometer races, and it maximizes the vertical rate of ascent for a given metabolic power (Giovanelli et. al. 2016). If we extrapolate our Equation 1 to 30°, it predicts a PTS of 0.983 m/s. Using the rule of thumb from Hubel and Usherwood (2013), a 1% decrease in PTS per 1% incline, with the baseline PTS at 0° found in the present study (1.95 m/s), the predicted PTS at 30° would be 1.09 m/s. There is currently no published research regarding the PTS at 30°. Extrapolating the EOTS to 30° using equation 2 yields a value of 0.909 m/s for the EOTS. From Ortiz et. al. (2017), this predicted EOTS closely corresponds to the data for the lone subject who had the aerobic fitness needed to reach EOTS at such a steep incline. Ultimately, these equations and predictions provide information on the relationship between PTS and EOTS on steeper inclines, which is often difficult to determine (Ortiz et. al. 2017).

Our study had some limitations. First, when trail/mountain runners walk on inclined terrain outdoors, they often place their hands on their quadriceps to facilitate knee extension during late stance. However, due to the constraints of the mouthpiece and breathing tube used to collect the expired air, subjects were unable to flex at the waist in order to place their hands on their knees during inclined walking. This may have influenced metabolic cost and discomfort in walking, especially at 10° and 15°, and thus slightly distorted the calculated EOTS.

A minor limitation occurred during our EOTS calculation. The goal was to measure energetic cost for at least one speed faster and at least one speed slower than the subjects’ EOTS. However, at 15°, three subjects were unable to complete the faster trials with an RER < 1.0. As a result, those three subjects’ EOTS regression analyses failed to capture the linear regression equation intersection point and some extrapolation (by ∼0.1 m/s) was required. This extrapolation may have increased the variability of the calculated EOTS at 15°.

Many interesting questions remain around the topic of human gait transition on inclines. For example, we should more explicitly explore the idea that energetics have a greater influence on PTS at higher exercise intensities. Considering the strong evidence for a neuromechanical trigger for the PTS during level locomotion, future inclined gait transition research should study the influence of biomechanical parameters. Determining if any joint kinetic or kinematic variables trigger the uphill PTS are logical future aspects to study. Likewise, pairing the kinetic and kinematic investigation with a neuromuscular component, measuring electromyography (EMG) activity of surface leg muscles, would be helpful in determining how biomechanical parameters influence gait transition speed (Whiting et. al. in press). Finally, performing more sophisticated biomechanical measurements concurrently with EMG measurements would allow for follow-up of Neptune and Sasaki’s findings that muscle force-length and force-velocity properties hinder the plantar flexors when walking at the PTS (Neptune and Sasaki 2005).

## CONCLUSION

Over a range of inclines, we found that the preferred (PTS), energetically optimal (EOTS) and heart rate optimal (HROTS) walk-run transitions speeds are slower on inclines. PTS is slower than EOTS up to 10° but they converge at 15°. EOTS is not accurately predicted by HROTS. Energetic, biomechanical, and neuromuscular factors all influence gait transitions and should be studied in further detail, especially on inclines commonly experienced by trail/ mountain runners, for whom the question of gait transition has large performance implications.

## LIST OF ABBREVIATIONS

bpm: beats per minute
EMG: electromyography
EOTS: Energetically Optimal Transition Speed
GPS: Global Positioning System
HROTS: Heart Rate Optimal Transition Speed
PTS: Preferred Transition Speed
RER: Respiratory Exchange Ratio
RWTS: Run-to-walk transition speed
WRTS: Walk-to-run transition speed

## ACKNOWLEDGMENTS

We thank: Robbie Courter and Derek Wright for their help with statistical analysis and results compilation respectively; Kyle Stearns, Clarissa Whiting, Ross Wilkinson, and Lance Perkins for help with data collection and all of the subjects for their willingness to participate.

## AUTHOR CONTRIBUTIONS

JB and RK - conception and design of research

JB - conducted experiments, analyzed data, statistics

JB and RK - interpreted results of experiments

JB - prepared figures

JB - drafted manuscript

JB and RK - edited and revised manuscript

JB and RK - approved the final version of the manuscript

